# A Sequel to Sanger: Amplicon Sequencing That Scales

**DOI:** 10.1101/191619

**Authors:** Paul DN Hebert, Thomas WA Braukmann, Sean WJ Prosser, Sujeevan Ratnasingham, Jeremy R deWaard, Natalia V Ivanova, Daniel H Janzen, Winnie Hallwachs, Suresh Naik, Jayme E Sones, Evgeny V Zakharov

## Abstract

Although high-throughput sequencers (HTS) have largely displaced their Sanger counterparts, the short read lengths and high error rates of most platforms constrain their utility for amplicon sequencing. The present study tests the capacity of single molecule, real-time (SMRT) sequencing implemented on the SEQUEL platform to overcome these limitations, employing 658 bp amplicons of the mitochondrial cytochrome *c* oxidase I gene as a model system. By examining templates from more than 5,000 species and 20,000 specimens, the performance of SMRT sequencing was tested with amplicons showing wide variation in GC composition and varied sequence attributes. SMRT and Sanger sequences were very similar, but SMRT sequencing provided more complete coverage, especially for amplicons with homopolymer tracts. Because it can characterize amplicon pools from 10,000 DNA extracts in a single run, the SEQUEL reduces costs 40-fold from Sanger analysis. Reflecting the capacity of each instrument to recover sequences from more than five million DNA extracts a year, this platform facilitates massive amplicon characterization.

## INTRODUCTION

High-throughput sequencers are doubling their analytical capacity every nine months (1,2), but their reads are generally short (<400 bp) and error rates reach 0.8%–1.7% (3). These limitations are an important constraint in three contexts: de novo genome assemblies are difficult (4), complex regions of well-known genomes can be intractable (5), and sequencing long amplicons is inefficient. Because of the latter constraint, Sanger sequencing is still widely used for amplicon characterization (6–9) despite its relatively high cost (10). While recent studies have established that Illumina (11,12) and Ion Torrent (10) platforms can analyze 1 kb amplicons with good accuracy, their need to concatenate short reads creates risks to data quality linked to the recovery of chimeras and pseudogenes. As well, because of their relatively complex workflows, costs are only three to four times less than those for Sanger analysis.

In contrast to the short reads delivered by other HTS platforms, the SEQUEL from Pacific Biosciences generates up to 60 kb reads (13,14). Despite its high error (13%) in single base calls (3), its long reads permit the generation of circular consensus sequences (CCSs) (15). For example, presuming a 12 kb read, each nucleotide position in a 1 kb amplicon is reanalyzed 10 times, allowing its accurate characterization. Since each run generates about 200,000 CCSs, the SEQUEL has the potential to analyze a diverse pool of amplicons. However, because each CCS reflects the characterization of a single molecule, SMRT analysis can recover heterogeneous sequences from a DNA extract, reflecting both variation in the target gene and diversity introduced by polymerase error. This contrasts with Sanger sequencing where the base call at each position integrates the signal from many amplicons so variants that comprise less than 10% of the amplicon pool have no impact on the base call at a particular position. Given this difference, empirical studies are needed to understand the complexities that arise when SMRT sequencing is employed to characterize amplicons.

A rigorous performance test demands the examination of amplicons with varied GC content and with substantial genetic divergence to reveal error biases dependent on sequence context (16). For example, long homopolymer runs are a challenge for Sanger sequencing (17,18), while Illumina platforms are subject to GC bias (16). Specimens employed to create DNA barcode reference libraries provide an ideal test system as Sanger reference sequences are available for a 648 bp region of the mitochondrial cytochrome *c* oxidase I (COI) gene from more than 500,000 species (19). Because of its considerable variation in GC composition (15–45%) and substantial sequence divergence, COI is sometimes challenging for Sanger analysis, usually as a result of homopolymer tracts. Consequently, this gene region provides a stringent test for the capacity of a sequencing platform to support amplicon analysis.

The present study tests the performance of SMRT sequencing in the characterization of COI amplicons from 20,000 DNA extracts, each from a different specimen. It compares the performance of SMRT and Sanger analysis in three key metrics: sequence length, sequence quality, and recovery success. It also ascertains the diversity of COI amplicons that can be characterized by a SMRT cell, the analytical chip for SEQUEL. Although this limit will depend on the number of reads and on the capacity to standardize amplicon abundances, the present study provides a first sense of the upper bound.

## MATERIALS AND METHODS

### Amplicon libraries

Four libraries were generated (Table 1) which collectively included amplicons from more than 20,000 specimens of Arthropoda, the most diverse animal phylum. These libraries spanned roughly three orders of magnitude in complexity as measured by the number (100–10,000) of different DNA extracts that were amplified to create templates that were pooled for analysis. Two low complexity libraries (#1–95 extracts, #2–948 extracts) were used to obtain high read coverage per extract, while the other two libraries (#3–9120 extracts, #4–9830 extracts) tested the upper limit on the number of samples that could be pooled.

**Table 1.** The geographic origin, number of specimens analyzed, and number of species represented in the four libraries analyzed by Sanger and SMRT sequencing.

| Library | Taxa | Collection Site | Specimens | Species | BOLD dataset |
| --- | --- | --- | --- | --- | --- |
| 1 | Insecta + Arachnida | Ontario, Canada | 95 | 95 | DS-PBBC1 |
| 2 | Insecta + Arachnida | Ontario, Canada | 948 | 945 | DS-PBBC2 |
| 3 | Insecta + Arachnida + Collembola | Para, Brazil & Selangor, Malaysia | 9,120 | 2837 | DS-PBBC3 |
| 4 | Insecta | Guanacaste, Costa Rica | 9,830 | 1840 | DS-PBBC4 |

The DNA extracts used to construct libraries #1/2 were selected from a study that recovered COI sequences from 44,000 Canadian arthropods representing 5,600 species (20). From this array, 948 specimens were selected for SMRT analysis. Library #1 included 95 taxa – a single specimen from 82 insect species (Coleoptera–18, Diptera–12, Hemiptera–13, Hymenoptera–24, Lepidoptera–15) and 13 arachnid species (Mesostigmata–9, Sarcoptiformes–1, Trombidiformes– 3). Library #2 included 948 specimens, all belonging to a different species with two exceptions: one dipteran was represented by three and one hymenopteran by two specimens to test the impact of replication on the number of reads for a species. The 945 species in library #2 included 837 from the five major insect orders (Coleoptera–159, Diptera–245, Hemiptera–94, Hymenoptera–244, Lepidoptera–95) and 108 from six orders of arachnids (Araneae–23, Mesostigmata–38, Opiliones–3, Pseudoscorpiones–1, Sarcoptiformes–3, Trombidiformes–40). Aside from ensuring taxonomic diversity, the primary criterion for the selection of a species was its possession of >6% sequence divergence at COI from any species already included in the library, barring four species pairs (2 dipterans, 1 hemipteran, 1 mite) with low divergence (1.6– 3.2%) that were included to verify that SMRT sequencing could discriminate them. The resultant array of species showed substantial variation (20.4–44.1%) in the GC content of their COI amplicons.

Libraries #3/4 were used to test the number of different amplicons that could be analyzed with a SMRT cell. Library #3 included amplicons from 9,120 arthropods from Brazil and Malaysia, while library #4 included amplicons from 9,830 arthropods from Costa Rica. Because some species in both libraries were represented by two or more specimens, the species count was lower than the sample size (4,764 species versus 18,950 specimens). The results from this analysis also permitted a comparison of the relative success of Sanger and SMRT sequencing in recovering COI from a diverse array of taxa.

### Molecular protocols

The same DNA extracts were employed for Sanger and SMRT sequencing (Figure 1). They were generated using a membrane-based protocol (21) which extracted DNA from a single leg of larger specimens or the whole body of smaller taxa (22). Particularly in the latter case, the amplicons from a particular DNA extract might derive from several sources because of the co-extraction of DNA from endosymbionts, parasitoids, and prey in the digestive tract.

**Figure 1.**
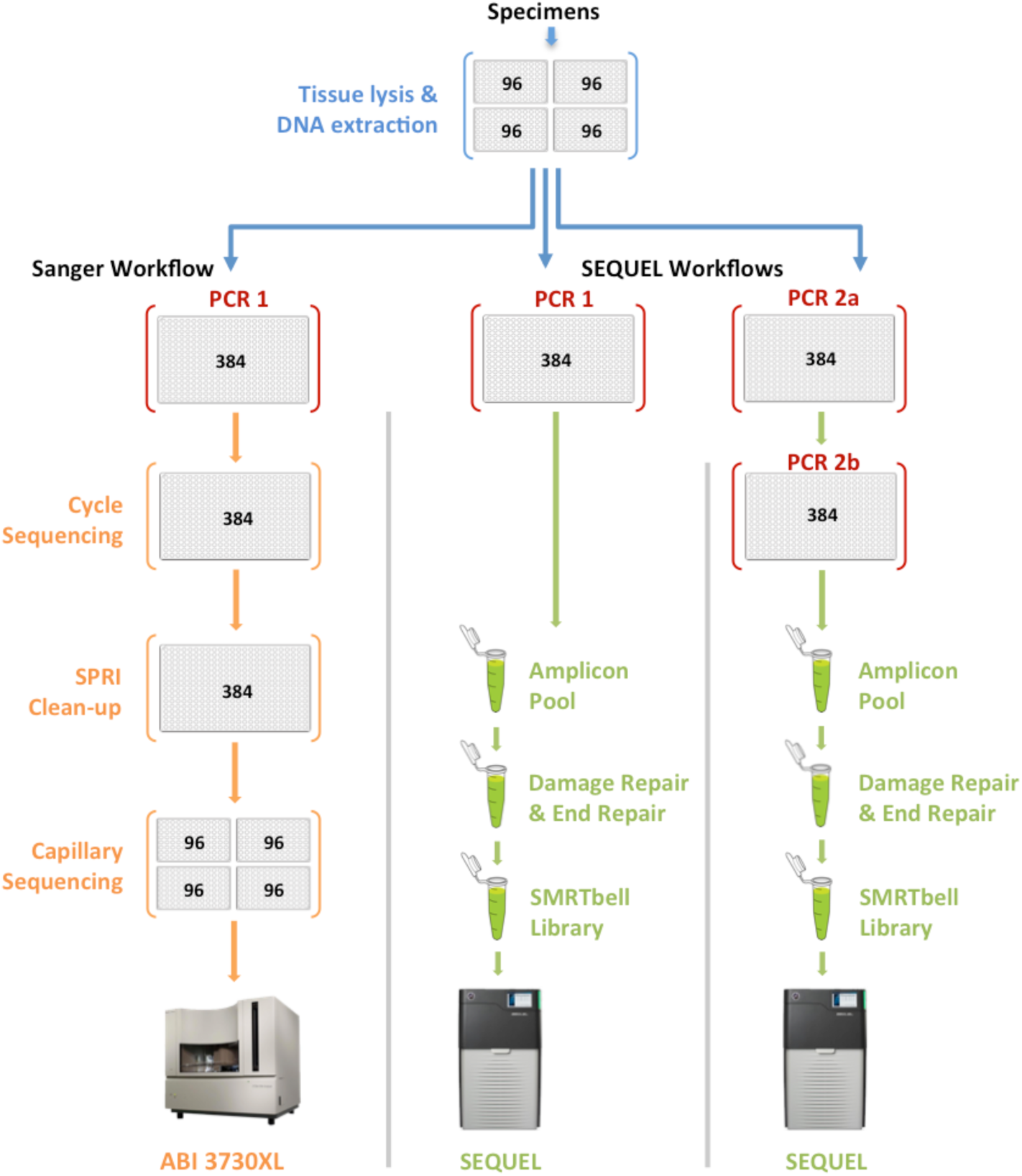
A comparison of the analytical pipelines for Sanger and SMRT sequencing. Blue arrows indicate shared steps in the workflow: tissue lysis and DNA extraction (plates marked in blue) and PCR (plates marked in red). Orange arrows indicate stages specific to the Sanger workflow while green arrows represent steps in the SMRT workflow.

Each DNA extract was used as a template for COI amplification without normalization of its concentration. Two PCR protocols, both targeting the same 658 bp segment of COI, were employed to generate amplicons for Sanger and SMRT sequencing. PCR1 was used when amplicon tagging was not required to link sequence records to their source, either because the amplicons were analyzed individually (i.e. Sanger sequencing) or because deep COI divergences among samples permitted *post hoc* taxonomic assignments (i.e. SMRT sequencing of libraries #1/2). PCR2 was used for SMRT sequencing of libraries #3/4 to enable the association of each CCS to its source well/specimen. Each PCR reaction was composed of 5% trehalose (Fluka Analytical), 1x Platinum Taq reaction buffer (Invitrogen), 2.5 mM MgCl__2__ (Invitrogen), 0.1 μM of each primer (Integrated DNA Technologies), 50 μM of each dNTP (KAPA Biosystems), 0.3 units of Platinum Taq (Invitrogen), 1 μL of DNA extract, and Hyclone ultra-pure water (Thermo Scientific) for a final volume of 6 μL.

PCR1 employed a single primer cocktail, C_LepFolF, C_LepFolR (23), and the following thermocycling protocol (initial denaturation for 2 min at 94°C, then 5 cycles of denaturation for 40 sec at 94°C followed by annealing for 40 sec at 45°C and extension for 1 min at 72°C, then 35 cycles of denaturation for 40 sec at 94°C followed by annealing for 40 sec at 51°C and extension for 1 min at 72°C, followed by final extension for 5 min at 72°C). Most reactions generated a 709 bp product (658 bp of COI plus 51 bp of forward and reverse primers), but some were slightly shorter because certain species had a 3–15 bp deletion in COI. To minimize the number of analytical steps, PCR amplicons were not purified and their concentrations were not normalized prior to sequence characterization, but the success of PCR was evaluated by scoring E-gels (Thermo Fisher) to confirm the presence of an amplification product. Products for Sanger sequencing were diluted 1:4 with ddH__2__O before 2 μl was used as template for a cycle sequencing reaction. As well, 2 μl from each of the 95/948 amplicons in libraries #1/2 were pooled to create two amplicon mixtures that were submitted for SMRT sequencing.

PCR2 involved two rounds of amplification (PCR2a, PCR2b). The first round (PCR2a) used a single primer cocktail (C_LepFolF, C_LepFolR) tailed with 30 bp adapter sequences (AF=gcagtcgaacatgtagctgactcaggtcac; AR=tggatcacttgtgcaagcatcacatcgta). These tails provided binding sites for primers tagged with unique molecular identifiers (UMIs) that were introduced in PCR2b (24,25). PCR2a employed the following thermocycling regime (initial denaturation for 2 min at 94°C, then 20 cycles of denaturation for 40 sec at 94°C followed by annealing for 1 min at 51°C and extension for 1 min at 72°C, followed by final extension for 5 min at 72°C). After a 1:2 dilution with ddH__2__O, the products were used as template for PCR2b which employed PCR primers consisting of a terminal 5 bp pad sequence (GGTAG), a 16 bp UMI, and a 30 bp AF or AR adapter to match the primer tails from PCR2a. Because the SEQUEL platform is well suited for asymmetric UMI tagging, 100 forward and 100 reverse primers, each with a different UMI, permitted 10,000 pairwise combinations, making it possible to attribute every sequence to its source by deploying a unique primer combination in each well. For example, discrimination of the 96 negative controls and 9,120 samples in library #3 required 96 UMI-F and 96 UMI-R primers to create 9,216 primer combinations. A Biomek FX liquid handler with a 384-channel head was employed to avoid errors in dispensing the designated primer cocktail into each well. PCR2b employed a thermocycling regime identical to PCR2a except the annealing temperature was raised to 64°C. Although the amplicons generated by PCR2b were ordinarily 811 bp long (10 bp pad, 32 bp UMIs, 60 bp AF/AR adaptors, 51 bp COI primers, 658 bp COI), some were 3– 15 bp shorter because of deletions in the COI gene itself. A 2 μl aliquot from each of the 9,216 PCRs for library #3 (9,120 samples, 96 negative controls) and from each of the 9,932 PCRs for library #4 (9,830 samples, 102 negative controls) were pooled to create two libraries for SMRT analysis.

### Sanger sequencing

Each product from PCR1 was Sanger sequenced using BigDye v3.1 from Life Technologies (Thermo Fisher). Sequencing reactions were performed by adding 0.5 μl of each diluted PCR product (95 for #1, 948 for #2, 9,216 for #3, 9,932 for #4) into 384-well plates prefilled with 5 μl of sequencing reaction mix following the manufacturer’s protocol. All cycle sequencing products were purified using an automated SPRI method (26) and sequenced on an ABI 3730XL. Libraries #1/2 were sequenced in both directions while libraries #3/4 were sequenced in one direction using the C_LepFolR primer. Trace files were submitted to the Barcode of Life Datasystems (BOLD; http://www.boldsystems.org) where they underwent quality trimming and filtering to produce barcode sequences that subsequently gained a Barcode Index Number (=species) assignment (27).

### SMRT sequencing

DNA quantity was evaluated for each amplicon pool using a Bioanalyzer and Nanodrop system before a 1 μg aliquot from each pool was used to prepare a SMRTbell library (28). Prior to ligation of the hairpin adapters that bind the sequencing primer and DNA polymerase, amplicons underwent damage- and end-repair to create double-stranded amplicon fragments with blunt ends. The resulting SMRTbell libraries were purified with AMPure® PB magnetic beads and combined with a sequencing primer and polymerase before each was loaded into a single SMRT cell to quantify amplicon diversity. Each of the resultant CCSs represented the analysis of a concatenated set of sub-reads, each corresponding to a single pass through a particular SMRTbell. Although the reads generated by each SMRT cell were output as a single fastq file, they varied in quality, reflecting, in part, variation in read length. Following convention, the number of CCS reads was determined for three data partitions (99%, 99.9%, 99.99%) where the percentage value indicates the proportion of bases in each CCS that is predicted to match its template based on Pacific Bioscience’s model of the circular sequencing process.

### Analysis of SMRT data

Slightly different analytical paths were required to analyze the results from libraries #1/2 versus #3/4 because SMRT sequences from the latter libraries included UMI tags. However, once each UMI-tagged CCS was assigned to its source well (details below), the four datasets were analyzed on mBRAVE (http://www.mbrave.net) using a standard pipeline which involved sequence trimming, quality filtering, de-replication, identification, and OTU generation. Because of its integration with BOLD (29), mBRAVE has direct access to the reference libraries needed for data interpretation.

A key step in data analysis involved the categorization of each CCS as ‘target’ or ‘non-target’. This assignment required pairwise comparisons against the reference Sanger sequences for each library (available as DS-PBBC1, DS-PBBC2, DS-PBBC3, DS-PBBC4 on BOLD). Prior to assignment, each CCS was trimmed by excising 30 bp from its 5’ and 3’ termini to ensure removal of the 25/26 bp primers and it was then further truncated to 648 bp. After trimming, the quality of each CCS was assessed; those with a mean QV <40, length <500 bp or >1% of their bases with a QV <20 were excluded. All remaining sequences were de-replicated based on perfect string identity and each distinct CCS was examined for similarity to the Sanger reference sequences in its library. In the case of libraries #1/2, each CCS with >98% similarity to any one of the 95/948 Sanger references was assigned to the ‘target’ category while those with lower similarity were ‘non-target’. The same sequence similarity value was employed for libraries #3/4, but it was implemented on a well-by-well basis. For each library, the pairwise distance value used to categorize each CCS employed k-mer searches followed by verification through global pairwise alignment using the Needleman-Wunsch algorithm (30) to the appropriate reference sequence array. Sequences were only assessed for a match when there was >80% overlap between the CCS and the Sanger reference, but less than 0.01% of all sequences were excluded based on this filter.

The fastq files for libraries #3/4 required an initial step to assign each CCS to its source well, work that was completed with a pipeline constructed from open source tools and python scripts. Pad sequences were trimmed from both ends of each CCS using *cutadapt* (31). To de-multiplex the fastq files, each CCS was first split at the UMI-F using a barcode splitter from the *fastx* toolkit (2017; http://hannonlab.cshl.edu/fastx_toolkit/index.html), and then split at the UMI-R. Because sequencing of a SMRTbell template can start from either forward or reverse strands of the amplicon, about 50% of the reads had to be reverse complemented prior to scoring the UMIs. Only CCSs with a perfect match to a UMI were retained. This stringent criterion meant that many CCSs (38.4% for #3, 40.7% for #4) were not assigned to a well. Most of these cases reflected erosion of the pad and the adjacent UMI tag that likely occurred during the damage- and end-repair stages of SMRTbell ligation. All sequences with a perfect UMI match were trimmed with a hard cut of 46 bp at both ends using the *fastx* trimmer to remove the UMI (16 bp) and the AF/AR adaptor (30 bp). This process produced sequence records for each COI amplicon and the primers employed in its PCR, each assigned to its source well by examining its UMI-F/UMI-R. In the usual situation where multiple sequence records were recovered from a well, their congruence was determined. Whenever the CCSs from a well possessed >2% divergence from any sequence in the reference library, they were assigned to a new molecular operational taxonomic unit (MOTU). Each MOTU was compared with all sequences on BOLD to ascertain if it derived from a bacterial endosymbiont such as *Wolbachia* or from another arthropod, reflecting contamination.

### Comparison of Sequence Recovery by Sanger and SEQUEL

All specimens in Libraries #3/4 were analyzed on both platforms, permitting comparison of the success in sequence recovery via Sanger and SMRT sequencing. Success in sequence recovery required meeting four criteria: 1) mean QV>35; 2) <1% of bases with QV<10; 3) <5% of bases with QV<20; and 4) minimum length = >493 bp (75% of the barcode region).

## RESULTS

### Sanger sequence metrics

The sequences for libraries #1/2 had greater mean QV scores (53.8 vs 51.8) and lengths (640 bp vs 569 bp) than those for libraries #3/4, reflecting the fact that they were obtained via bidirectional analysis while the latter libraries were unidirectionally sequenced (Figure 2). The results showed that 9% of the reads for libraries #3/4 failed to meet the four quality criteria set for recognition as successful sequence recovery.

**Figure 2.**
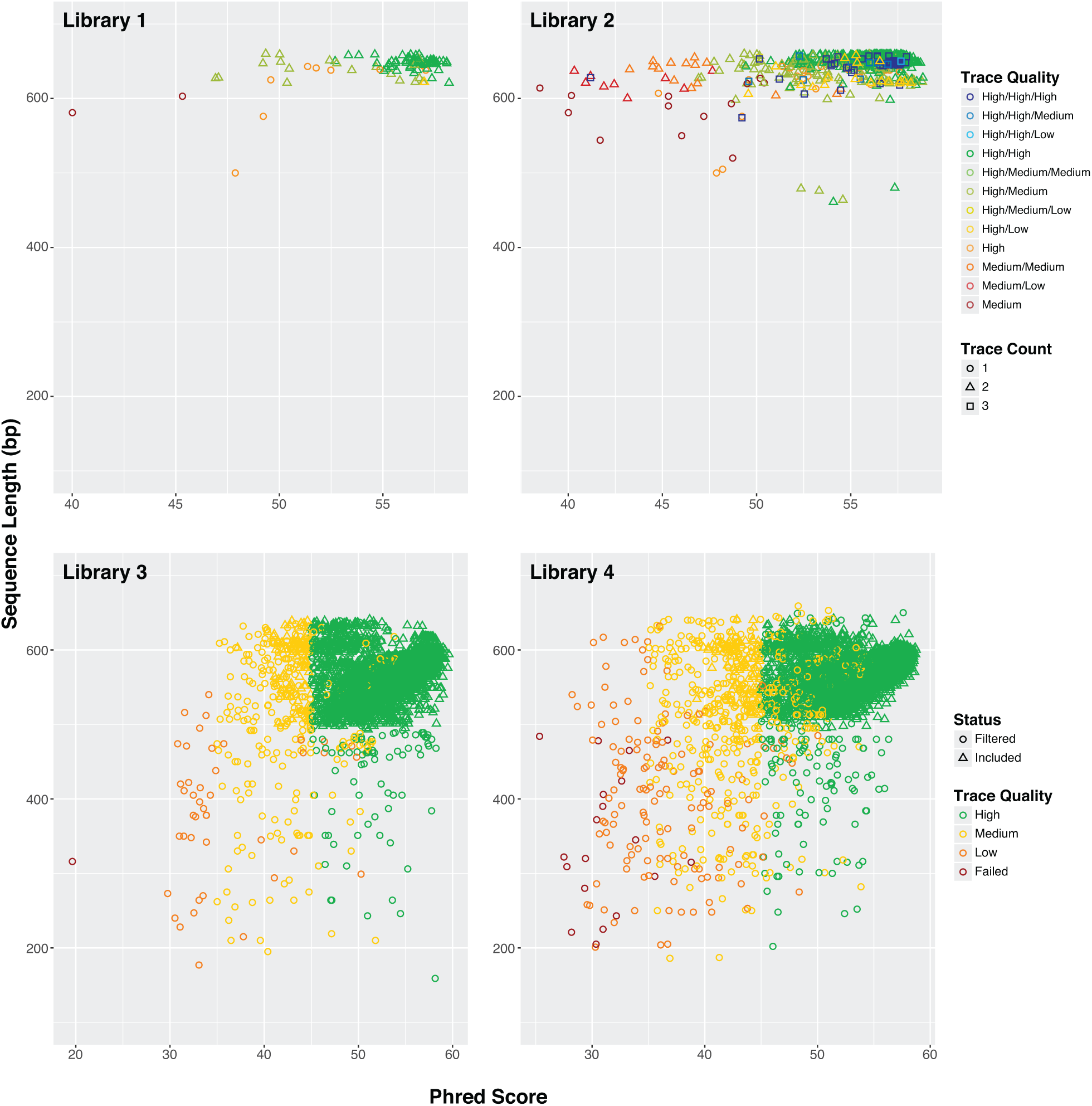
Variation in the quality and length of Sanger reads for the four libraries. Libraries #1/2 were sequenced bidirectionally while libraries #3/4 were sequenced unidirectionally. Three trace files were generated for a few difficult amplicons in library #2 to maximize succes in sequence recovery.

### SMRT sequence metrics

The number of reads per SMRT cell averaged 470,000 with a mean length of 13.8 kb, but about half failed to qualify for CCS analysis (Table 2). The number of CCSs varied less than two-fold among the four runs with an average of 244,000 in the 99% data partition versus 167,000 in the 99.9% partition and 64,000 in the 99.99% partition. A higher proportion of the CCSs for libraries #1/2 than #3/4 qualified for the 99.99% partition, presumably because their amplicons were 14% shorter (709 bp versus 811 bp).

**Table 2.**
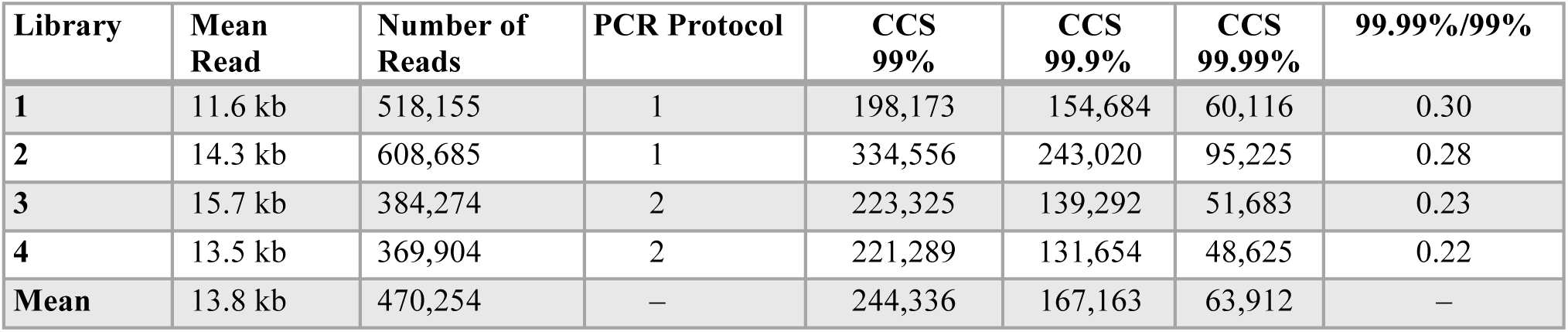
Mean read length and number of circular consensus sequences (CCSs) for COI from the four libraries in three data partitions.

| Library | Mean Read | Number of Reads | PCR Protocol | CCS 99% | CCS 99.9% | CCS 99.99% | 99.99%/99% |
| --- | --- | --- | --- | --- | --- | --- | --- |
| 1 | 11.6 kb | 518,155 | 1 | 198,173 | 154,684 | 60,116 | 0.30 |
| 2 | 14.3 kb | 608,685 | 1 | 334,556 | 243,020 | 95,225 | 0.28 |
| 3 | 15.7 kb | 384,274 | 2 | 223,325 | 139,292 | 51,683 | 0.23 |
| 4 | 13.5 kb | 369,904 | 2 | 221,289 | 131,654 | 48,625 | 0.22 |
| Mean | 13.8 kb | 470,254 | – | 244,336 | 167,163 | 63,912 | – |

While sequence quality was highest in the 99.99% partition, the difference among partitions was small (Figure 3). For example, QV scores only increased from 88 to 92.5, and mean sequence similarity to the Sanger references was above 99.8% in all three partitions. There was a sharper difference in the incidence of indels; they were 2–3 times more frequent in the 99% than the 99.99% partition. However, because indels averaged less than 1.5 per sequence in all partitions, they were readily recognized and excised following alignment. Because the quality differences were small and just 26% of the data in the 99% partition qualified for inclusion in the 99.99% partition, most subsequent analyses employ the 99.9% partition because it coupled high sequence quality with the retention of most (68%) of the CCS reads.

**Figure 3.**
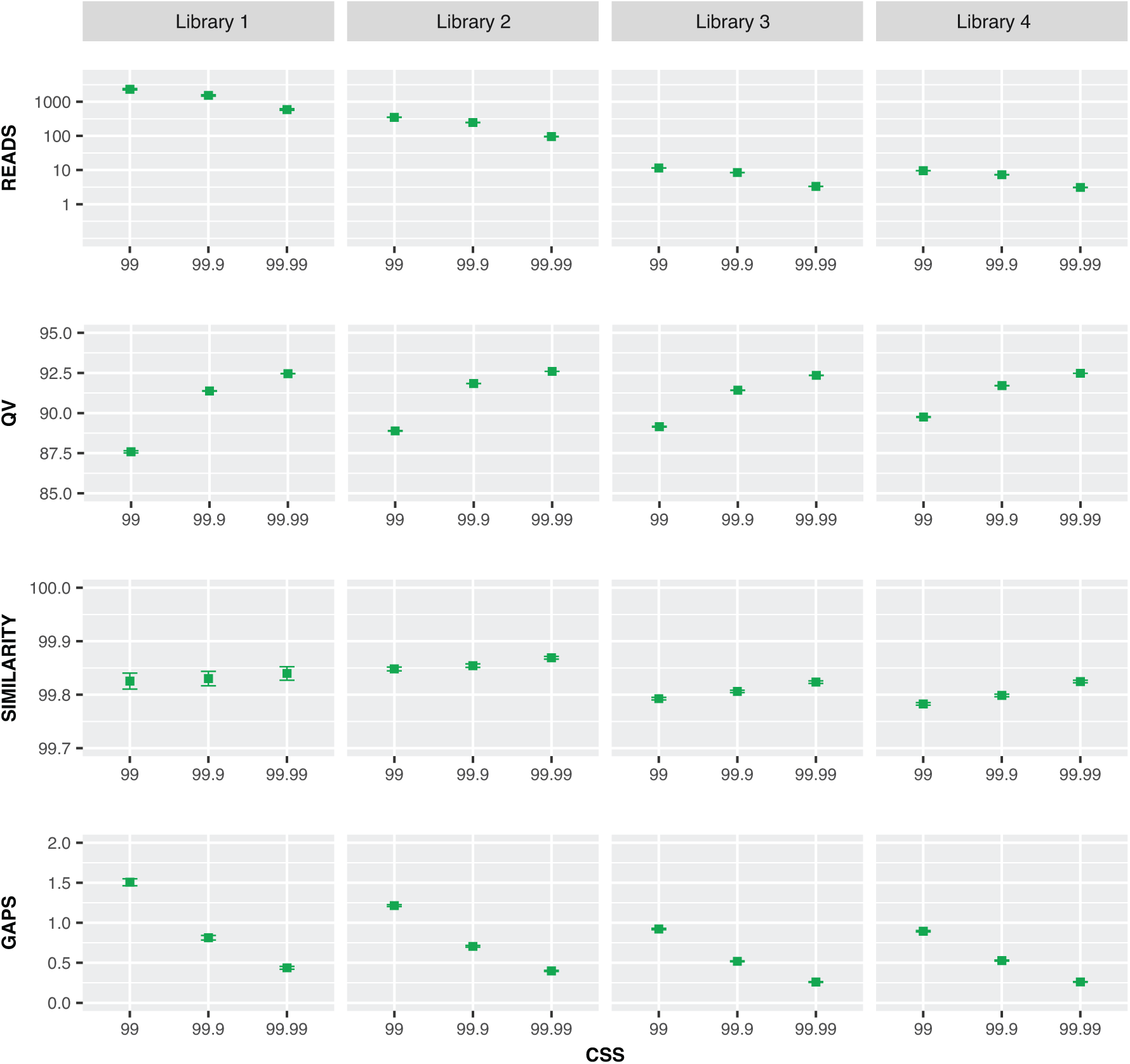
Mean +/-SE for four metrics (reads per DNA extract, QV scores for each CCS, CCS similarity to Sanger reference, indels per CCS) for three CCS partitions (99%, 99.9%, 99.99%) for the four COI libraries.

### Comparison of Sanger and SMRT sequences

#### Sequence Context

Examination of the COI sequences for the 945 taxa in Library #2 using both Sanger and SMRT sequences revealed five homopolymer tracts (Figure 4A, Figure S1). They ranged in length from 6 bp to 11 bp, but only one often showed >7 bp runs. Most of the >7 bp runs were thymine (95%) or cytosine (4%) (Figure 4B). Aside from this variation in nucleotide composition within amplicons, there were large differences among the 945 taxa in overall GC composition (15.2–41.6%).

**Figure 4.**
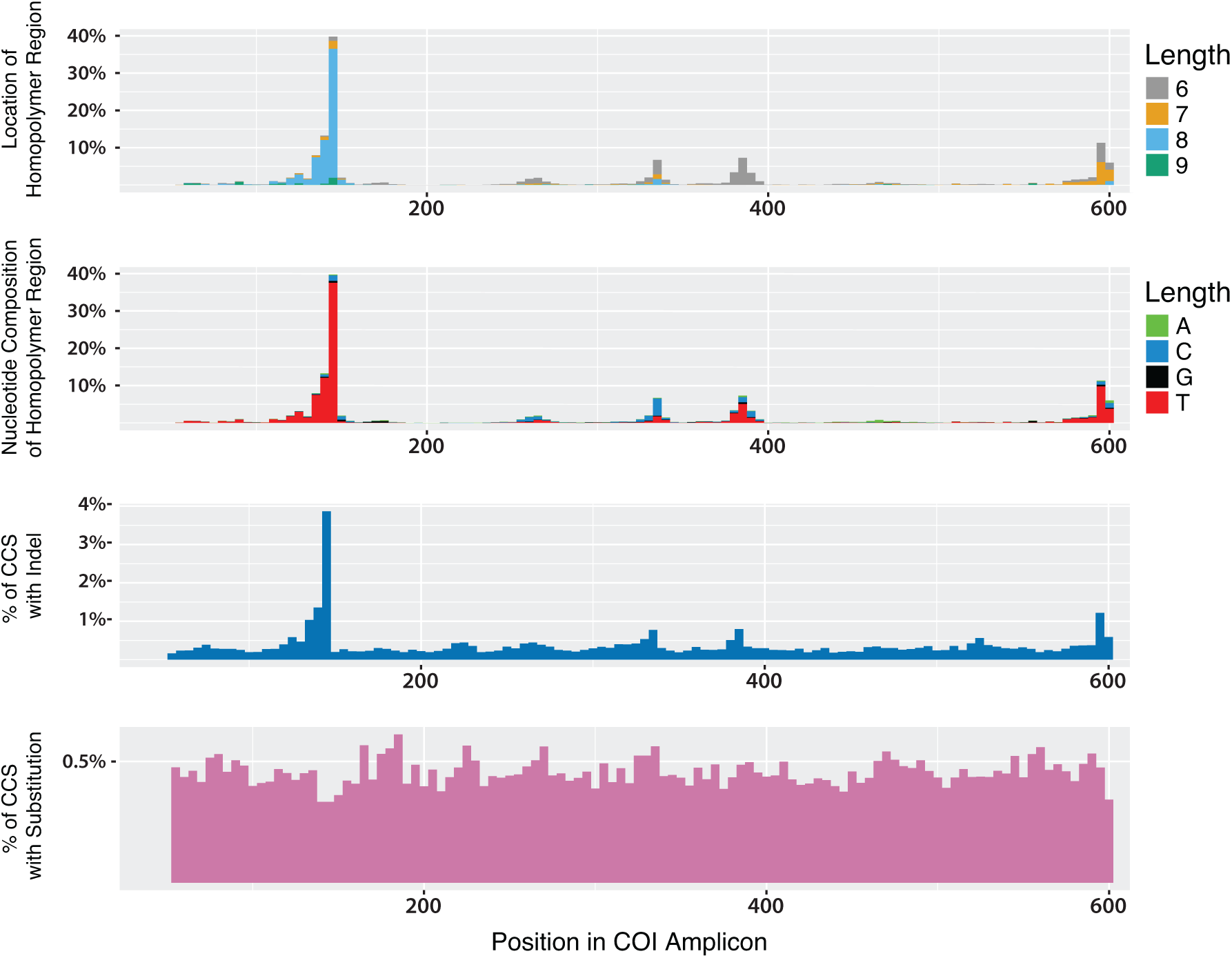
Sliding window (5 bp) analysis showing (**A**) the distribution and (**B**) the GC composition of the homopolymer regions in the COI gene for the 945 taxa in library #2. (**C**) The frequency of indels and (**D**) the frequency of substitutions in each window for the 99.9% SMRT partition. The incidence of substitutions and indels are shown per base pair.

#### Indels

A sliding window analysis of the SMRT sequences indicated that indels were generally infrequent, averaging 0.1% per base pair (Figure 4C), but their incidence rose 30-fold in the COI segment with the long homopolymer tract. The number of indels was not linked to the overall GC content of a COI sequence (Figure S2), but there was evidence for consistent differences among taxa as evidenced by the strong correlation in indel counts for the 95 taxa that were shared by libraries #1/2 (Figure S3).

#### Substitutions

A sliding window analysis of the SMRT sequences indicated that the frequency of nucleotide substitutions was nearly stable across the amplicon, averaging 0.5% per base pair. As a result, the SMRT and Sanger sequences for a particular specimen showed close concordance. For example, all 95 species in library #1 showed <0.3% divergence between their Sanger and SMRT sequences (Figure 4D). A NJ tree for the 945 taxa in library #2 also demonstrated a close correspondence between Sanger and SMRT sequences (Figure S4) and further indicated the clear separation of the four species pairs with low sequence divergences (1.67%, 1.85%, 2.00%, 3.20%) The same pattern was evident for Libraries #3/4 although this was demonstrated more compactly by plotting sequence divergences between the sequences generated by Sanger and SEQUEL analysis (Figure S5).

#### Sequence lengths

Sanger sequences averaged 71 bp longer for libraries #1/2 than #3/4 because amplicons from the first two libraries were sequenced bidirectionally while the latter were only analyzed unidirectionally (Figure 5). The SMRT sequences showed considerably less length variation than even bidirectional Sanger reads. In fact, most SMRT sequences shorter than 658 bp involved taxa with a deletion in the COI gene itself.

**Figure 5.**
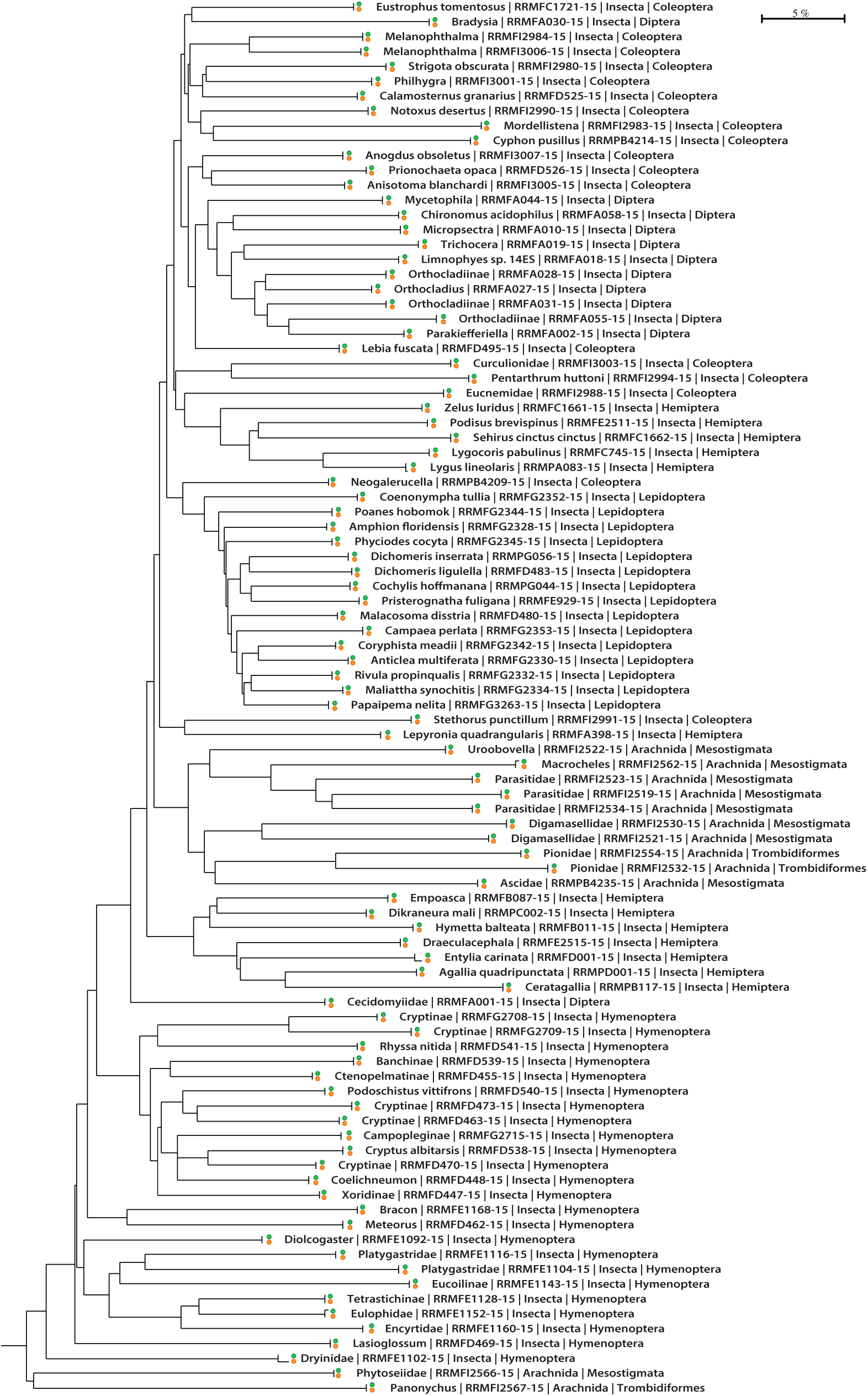
Neighbor-Joining tree showing the correspondence between Sanger (orange) and SMRT (green) COI sequences for the 95 taxa in library #1.

### Sequence recovery via Sanger and SMRT analysis

All specimens in libraries #1/2 possessed a Sanger sequence as this was a requirement for their inclusion, but the present study established that SMRT analysis was highly effective in their recovery from a multiplexed sample. (Figure 6). In fact, all three SMRT partitions recovered the 95 taxa in library #1 while the 99% and 99.9% partitions recovered the 945 taxa in library #2. Despite its considerably lower CCS count, the 99.99% partition only lacked coverage for two of the taxa in library #2.

**Figure 6.**
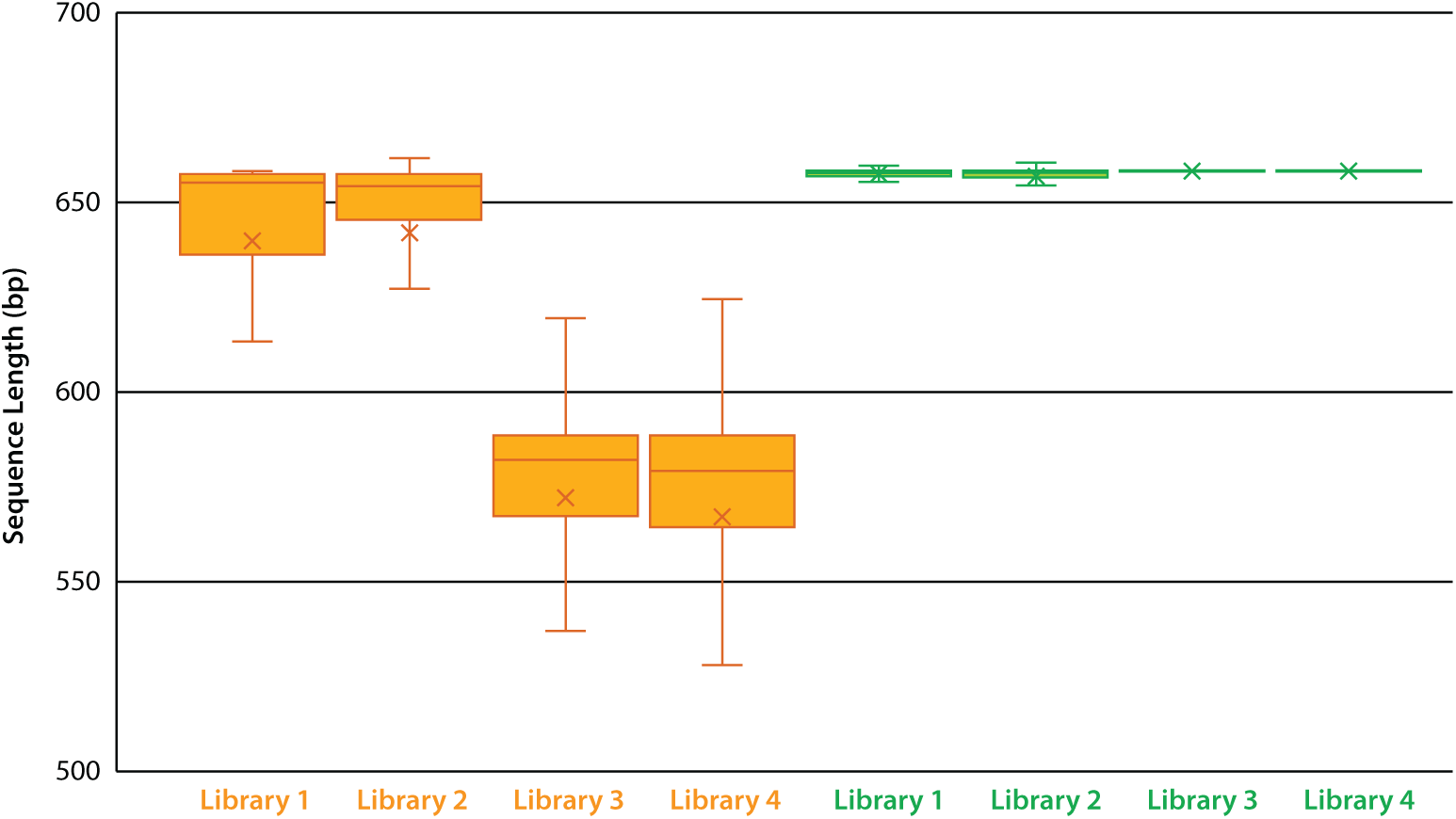
Box-plot showing mean length of sequences recovered from four COI amplicon libraries via Sanger (orange) and SMRT (green) analysis.

Sanger analysis produced a sequence from 8134 of the specimens in library #3 (89.2%) and from 8207 of those in library #4 (83.5%). SMRT analysis recovered sequences from more specimens in both the 99% and 99.9% partitions (90.0%, 85.8% for latter partition), but fewer for the 99.99%. Some specimens in library #3 (1.1%) and #4 (1.6%) failed to deliver a sequence with both Sanger and SMRT analysis, likely reflecting cases where primer binding failed. Further analysis of the results for libraries #3/4 indicated significant variation (X^2^ = 30.48, p<0.001) in Sanger sequence recovery among insect orders with 10–15% lower success for Coleoptera, Hemiptera, and Hymenoptera than for Diptera and Lepidoptera (Figure 7). SMRT analysis improved sequence recovery for all orders, particularly for Hymenoptera, reflecting its capacity to sequence amplicons with long homopolymer tracts.

**Figure 7.**
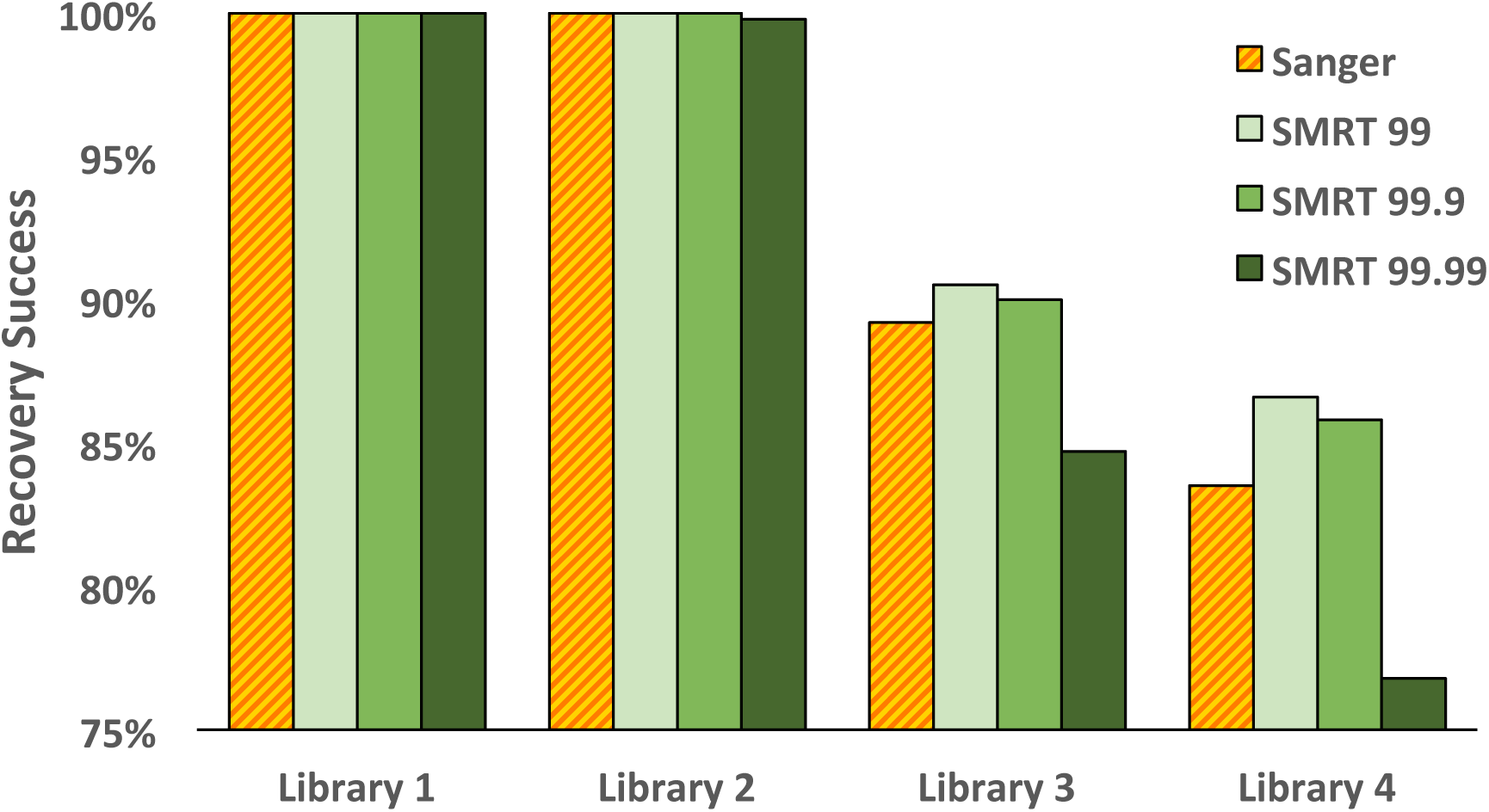
Success in the recovery of a COI sequence from the four libraries using Sanger and SMRT analysis. Recovery success with SMRT analysis is shown for three data partitions.

### Screening amplicon libraries with SMRT sequencing

CCS counts for each library approximated a normal distribution although those for libraries #3/4 were truncated (Figure 8). The CCS count per taxon declined with rising library complexity from a mean of 1515 for library #1 to 8.9/7.8 for libraries #3/4. The actual CCS count for the latter two libraries averaged 15, but many records could not be assigned to a source specimen because their UMI was eroded. The coefficient of variation in CCS counts was low, ranging from 31% for library #2 to 45% for #4, meaning there was only two-fold variation in the CCS count for about two thirds of the taxa in each library. Further evidence for the limited variation in CCS counts among specimens was provided by the two species in library #2 that were represented by more than one specimen. The dipteran with three individuals had the highest CCS count (1018 reads), while the hymenopteran with two individuals was in fifth place (487 reads) among the 945 species. CCS counts for the taxa in libraries #1/2 were positively correlated with the GC content of their amplicons, but libraries #3/4 did not show this association (Figure S6).

**Figure 8.**
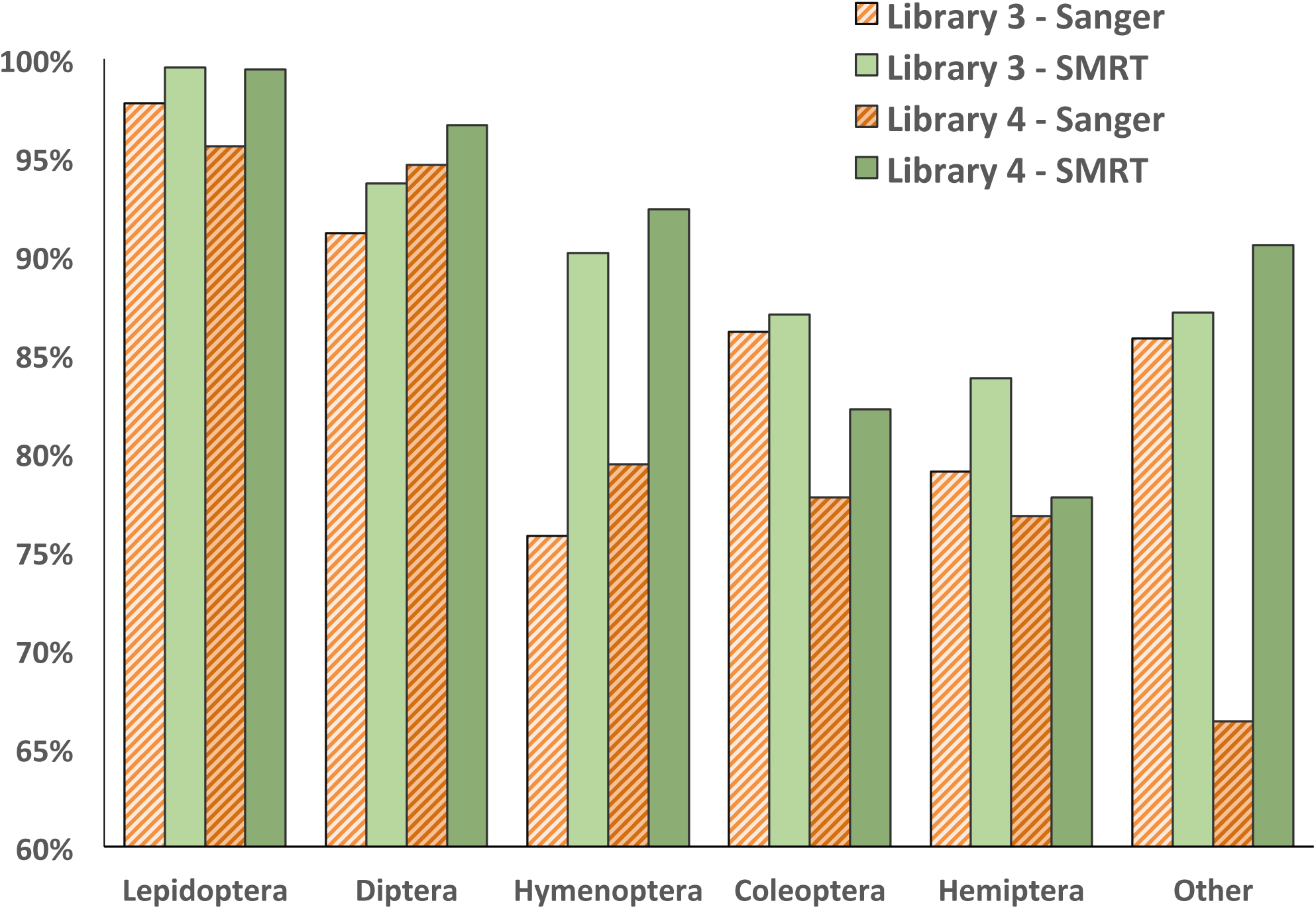
Comparison of the success in recovering >500 bp of COI from 4677 insect species belonging to varied orders by Sanger and SMRT sequencing.

**Figure 9.**
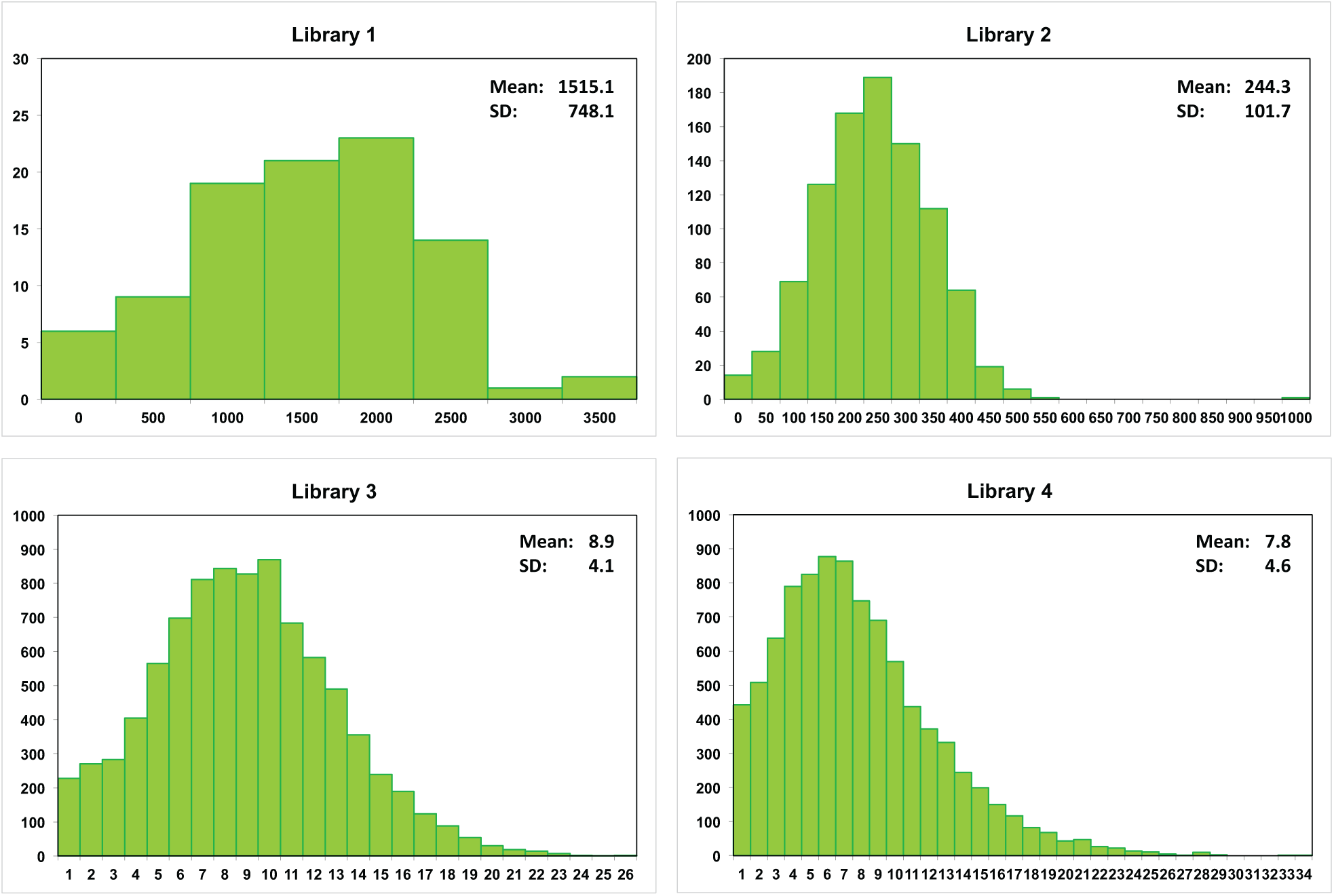
Number of CCS counts for COI in the 99.9% partition for each taxon in four libraries that were characterized with a single SMRT cell.

### Incidence of non-target sequences with SMRT analysis

Few sequences derived from the negative control wells in libraries #3/4. In fact, just two sequences were recovered from the 96 controls in #3, and five from the 102 controls in #4, each involving a single CCS per well. More non-target sequences were recovered from wells with a DNA extract. They comprised 2.2% of the sequences from libraries #1/2, although their source could not be identified because the samples were not UMI-tagged. Non-targets were slightly more abundant for libraries #3/4, reaching 3.1% and 4.0% respectively with most deriving from arthropods (84.4% in #3, 91.9% in #4) or endosymbiotic bacteria (14.8% in #3, 7.7% in #4). Non-target sequences were only present in higher abundance than the target sequence in 1.6% of the wells in library #3 (149/9210) and in 1.5% of those in #4 (193/9830). The mean CCS count for wells where non-targets predominated was significantly lower than those where the target was dominant (3.5 versus 8.9 reads for #3, 2.9 versus 7.8 reads for #4), and these wells often possessed low quality Sanger reads (Figure S7).

## DISCUSSION

The present study has shown that the SEQUEL is a highly effective platform for amplicon sequencing. Although the number of CCSs generated by each SMRT cell was modest (167,000 in the 99.9% partition), these sequences showed close congruence to their Sanger counterparts. Earlier studies have reported similar results, but they focused on 16S rRNA and examined far fewer templates (32–33). The present analysis sequenced amplicons from more than 5,000 species, taxa whose COI amplicons showed wide variation in GC content (15–45%) and diverse compositional attributes. Across this range of templates, SMRT sequencing showed no points of failure. SMRT sequences also had a major advantage over their Sanger counterparts as they regularly provided complete coverage for the target amplicon. By comparison, the generation of complete coverage through Sanger sequencing demanded bidirectional analysis to escape interpretational complexities introduced by ‘dye blobs’ and uncertainty in base calling at the initiation of each sequence (34). As a consequence of these factors, unidirectional Sanger reads were invariably truncated and even bidirectional reads showed more length variation than SMRT sequences, often reflecting barriers created by homopolymers.

Although single CCS records from a particular DNA extract corresponded closely with their Sanger counterpart, they were often not identical to it, averaging 0.6% divergence reflecting the presence of about three substitutions and 0.75 indels. Nucleotide substitutions occurred at similar frequency at each position in the COI amplicon, but indels showed site-specific variation, increasing 30-fold in regions with homopolymer tracts. The SMRT sequences recovered from such taxa typically showed several length variants, likely reflecting polymerase slippage in the homopolymer region during PCR (35). Because of its protein-coding function, SMRT sequences for COI could be readily aligned, allowing the excision of indels created through polymerase slippage. By comparison, the Sanger traces from taxa with long homopolymer runs were often uninterpretable. Viewed from this perspective, homopolymers disrupt Sanger analysis (18,35), but only lead to readily resolved length variation in SMRT sequences.

If the sequence variation noted among the CCS reads from individual DNA extracts derived from heteroplasmy in its source organism, one would expect a small number of variants (36). Instead, there was usually a single dominant sequence and many low-frequency variants with an indel and/or a substitution at varied positions, a pattern consistent with PCR errors (37). Presuming COI homoplasy in the source specimen, the frequency of errors in the final amplicon pool can be predicted from polymerase fidelity, sequence length, and the number of PCR cycles (38). As all reactions employed Platinum Taq and 40 cycles of amplification, 60% of the final amplicons should possess a PCR error, while 40% should match the source. This theoretical expectation coincided with the observed results; the dominant sequence represented approximately 40% of the sequences from each DNA extract, and perfectly matched its Sanger counterpart. The adoption of high fidelity DNA polymerases (39–40) could greatly diminish PCR errors, but their high cost and requirement for additional cleanup steps mean their use will only be justified in limited contexts.

This study has established that simple PCR protocols allow sufficient standardization of amplicon concentrations to permit a single SMRT cell to recover sequences from nearly 10,000 DNA extracts. Although the mean number of reads per taxon declined by more than three orders of magnitude (1515 to 8 as library complexity increased from 95 to 9830 templates), success in sequence recovery remained high (100% for 95 and 945, 90% for 9120, 86% for 9830). Because SMRT sequencing supports asymmetric UMI tagging, high levels of multiplexing could be implemented cost-effectively since just 200 primers were required to discriminate 10,000 samples (versus 50 times that many with symmetric tagging). Despite the modest number of reads generated by each run, SMRT sequencing supports high levels of multiplexing because the fidelity of each sequence is high. Although some samples failed to gain a sequence, the cost ($0.15) per sample was least at the highest level of multiplexing employed in this study and it could be further reduced. For example, the analysis of a library with amplicons from 40,000 DNA extracts on a single SMRT cell should generate an average of two reads and recover sequences from about 25,000 of them. Extracts without a sequence in the first run could then be pooled for secondary analysis, a strategy that would reduce sequencing costs to $0.05 per specimen. If deployed in all combinations, the 384 UMI tags currently available (https://github.com/PacificBiosciences/Bioinformatics-Training/wiki/Barcoding) can discriminate 73,536 amplicons, a capacity that will be useful once SMRT cells generate more CCSs. The high fidelity of SMRT sequencing has the additional advantage of minimizing the risk that UMIs will be misread, an error that leads sequences to be assigned to the incorrect source, a frequent problem with HTS platforms (41–42).

Prior studies have shown that other HTS platforms can sequence circa 1 kb amplicons, but their workflows are more complex and more expensive. For example, Shokralla et al. (43,11) demonstrated that the Illumina MiSeq could recover 648 bp COI amplicons from hundreds of taxa at a time. However, because it only generates 300 bp sequences (or 500 bp with its paired end approach), workflows required the acquisition of multiple reads of several amplicons to obtain a high-fidelity 648 bp sequence. As a consequence, the cost ($1.50) per extract is ten times higher than with SMRT analysis. Diekstra et al. (10) employed the Ion Torrent PGM platform to characterize up to 900 bp templates, but the amplicons had to be sheared to <300 bp before sequencing and reassembly. The resulting sequences corresponded closely with their Sanger counterparts, but analytical costs ($1.96) were only three-fold less than those for Sanger analysis ($6.00). Illumina and Ion platforms also run the risk of recovering chimeras when short reads are assembled to characterize longer amplicons. This is a particular risk for studies on mitochondrial genes as NUMTs are prevalent (44) and are readily recovered by PCR (45). Because most NUMTs are <300 bp (46), any strategy which relies on the amplification and assembly of short amplicons creates the risk of generating chimeras which combine a segment of the authentic gene with a NUMT. Perhaps reflecting this fact, Craud et al. (12) found that 5% of the COI sequences generated by MiSeq analysis diverged from their Sanger reference.

While Sanger analysis is effective for sequencing amplicons up to 1 kb (47,14), the characterization of longer templates requires primer walking or shotgun sequencing, protocols that introduce complexities and raise costs (48,49). The capacity of SMRT sequencing to analyze long amplicons is a key advantage in such situations (50). For example, the 5 kb CAD gene is valuable for phylogenetic studies, but its recovery via Sanger sequencing requires the analysis of six overlapping amplicons (51), and difficulties are often encountered in recovering the full set. Similarly, Zhang et al. (52) identified 1083 genes with high potential for phylogenetic studies on plants, but many exceeded 1500 bp in length. Because the average gene length across animals, fungi, plants, and protists ranges from 1200–1500 bp and from 1700–9500 bp when introns are included (53,54), the capacity of SMRT sequencing to analyze long amplicons is a general asset.

In summary, this study indicates that SMRT analysis is a powerful approach for amplicon sequencing. It can characterize templates with large divergence in GC content and long homopolymer tracts. SMRT sequences are congruent with those obtained through Sanger analysis, but analytical costs are reduced 40-fold. Because SMRT analysis supports massive multiplexing, a single SEQUEL platform can characterize millions of DNA extracts annually. Exploitation of this capacity is aided by the fact that data processing is simple. While Sanger sequencing requires the visual inspection of trace files to optimize data quality, SMRT sequences can be processed with an automated pipeline that is much simpler than those required for similar analyses on short-read HTS platforms.

## AVAILABILITY

Details on the specimens in each library coupled with Sanger sequences and trace files are available in four datasets on BOLD: library #1– http://dx.doi.org/10.5883/DS-PBBC1; library #2 – http://dx.doi.org/10.5883/DS-PBBC2, library #3 – http://dx.doi.org/10.5883/DS-PBBC3, library #4 – http://dx.doi.org/10.5883/DS-PBBC4. The 12 datasets holding the CCS records for each partition (99%, 99.9%, 99.99%) from libraries #1 to #4 are available in DRYAD under BioProject XXXXXX.

## SUPPLEMENTARY DATA

Supplementary Data are available at NAR online.

## ACKNOWLEDGMENT

We thank staff at Pacific Biosciences, particularly Jonas Korlach and Cheryl Heiner for aiding data acquisition. We are also very grateful to John Wilson for providing specimens from Malaysia and to Marlucia Martins from providing those from Brazil. We thank Suzanne Bateson for her aid with graphics.

